# Superficial white matter across the lifespan: volume, thickness, change, and relationship with cortical features

**DOI:** 10.1101/2022.07.20.500818

**Authors:** Kurt G Schilling, Derek Archer, Francois Rheault, Ilwoo Lyu, Yuankai Huo, Leon Y Cai, Silvia A Bunge, Kevin S Weiner, John C Gore, Adam W Anderson, Bennett A Landman

## Abstract

Superficial white matter (SWM) represents a significantly understudied part of the human brain, despite comprising a large portion of brain volume and making up a majority of cortical structural connections. Using multiple, high-quality, datasets with large sample sizes (N=2421, age range 5-100) in combination with methodological advances in tractography, we quantified features of SWM volume and thickness across the brain and across the lifespan. We address four questions: (1) How does U-fiber volume change with age? (2) What does U-fiber thickness look like across the brain? (3) How does SWM thickness change with age? (4) Are there relationships between SWM thickness and cortical features? Our main findings are that (1) SWM volume shows unique volumetric trajectories with age that are distinct from gray matter and other white matter trajectories; (2) SWM thickness varies across the brain, with patterns robust across individuals and across the population at the region-level and vertex-level; (3) SWM shows nonlinear changes across the lifespan that vary across regions; and (4) SWM thickness is associated with cortical thickness and curvature. For the first time, we show that SWM volume follows a similar trend as overall white matter volume, peaking at a similar time in adolescence, leveling off throughout adulthood, and decreasing with age thereafter. Notably, the relative fraction of total brain volume of SWM continuously increases with age, and consequently takes up a larger proportion of total white matter volume, unlike the other tissue types that decrease with respect to total brain volume. This study represents the first characterization of SWM features across the lifespan and provides the background for characterizing normal aging and insight into the mechanisms associated with SWM development and decline.

## Introduction

Superficial white matter (SWM) is the layer of white matter immediately beneath the cortex, and is composed of short-range association U-shaped fibers, or U-fibers, that primarily connect adjacent gyri (Guevara et al., 2020). As summarized in (Kirilina et al., 2020), these connections are unique, as they occupy a majority of the total white matter volume (Schüz et al., 2002), account for a majority of the connections of the human brain (Schüz et al., 2002; Schüz and Miller, 2002), are among the last parts of the brain to myelinate (Barkovich, 2000; Maricich et al., 2007; Schuz et al., 2006; Wu et al., 2014), and contain a relatively high density of neurons relative to other white matter systems (Suarez-Sola et al., 2009). Further, SWM has been shown to play a critical role in brain function, cognition, and disease (Guevara et al., 2020; Kirilina et al., 2020).

Despite its significance, SWM has been critically understudied compared to the long-range projection, association, and commissural fibers of the brain. This is largely attributed to challenges and limitations of diffusion MRI fiber tractography (Guevara et al., 2020; Schilling et al., 2018; Shastin et al., 2021), which is the primary modality used to study the structural connections of the human brain in vivo (Jeurissen et al., 2019). These challenges are associated with the complex anatomy of the SWM, including complex fiber orientation, fiber trajectories with high curvature, and partial volume effects with both gray matter and other white matter systems (Reveley et al., 2015; Shastin et al., 2021). However, recent advances in MRI data acquisition and image processing have made it possible to reliably study U-fibers in health and disease. This is typically done by measuring diffusion values such as fractional anisotropy (FA), mean diffusivity (MD), axial diffusivity (AD), and/or radial diffusivity (RD), and quantifying their changes in pathology. For example, decreases in anisotropy and increases in diffusivity of SWM have been documented in Alzheimer’s disease, autism spectrum disorder, schizophrenia, and healthy aging (Bigham et al., 2022; Bigham et al., 2020; Ji et al., 2019; Phillips et al., 2016; Reginold et al., 2016; Veale et al., 2021; Wu et al., 2016; Wu et al., 2014), among others (Guevara et al., 2020). These findings have been attributed to biological changes such as decreased coherence, decreased axonal packing, and thinning myelin, and further emphasize that SWM is especially affected in various pathologies.

Despite these developments, much is still unknown and uncharacterized about the SWM. For example, features such as SWM volume or thickness of the SWM sheet below the cortex have not been characterized. Unlike the *cortical thickness* and *cortical volume* metrics that have been thoroughly investigated, are easily quantified using standard toolsets (Destrieux et al., 2010; Fischl, 2012), and have proven strong associations with cognition, aging, and disease (Dickie et al., 2020; Dominguez et al., 2021; Fjell and Walhovd, 2010; Frangou et al., 2022; Gao et al., 2018; Habeck et al., 2020; Hou et al., 2021; Mattsson et al., 2018; Meyer et al., 2019; Racine et al., 2018; Roe et al., 2021; Steffener, 2021; Storsve et al., 2016; Tustison et al., 2019; Vogt et al., 2019), *SWM thickness* and *SWM volume* have not been investigated or quantified. Towards this end, we study three high-quality datasets that span the human lifespan (ages 5-100, N=2421 subjects), and implement advances and innovations in SWM tractography. First, we quantify total U-fiber volume and characterize how this tissue volume changes with age, making comparisons with total brain volume and total white and gray matter tissue volumes. Second, we quantify U-fiber thickness across the brain, and again describe how this changes with age in different areas of the brain. Finally, to investigate potential relationships between SWM and neighboring cortex, we explore correlations between SWM features and key features of the overlaying gray matter, including thickness, curvature, and sulcal depth. Together, this effort represents the first characterization of these features of the SWM and how they change across the lifespan and provides the background for characterizing normal aging and insight into the mechanisms associated with SWM development and decline.

## Methods

This study used three open-sourced datasets that span a large range of the human lifespan (*Data*), and advances in fiber tractography (*Superficial White Matter Tractography*) to map features of SWM volume and thickness across the brain (*Features and Analysis*). With these data, we aimed to answer four novel questions: (1) What does U-fiber thickness look like across the brain? (2) How does U-fiber volume change with age? (3) How does U-fiber thickness change with age? (4) Is there a relationship between U-fiber thickness and key cortical features: specifically, thickness, curvature, and sulcal depth?

### Data

The data used in this study come from the Human Connectome Project (Van Essen et al., 2012), which aims to map the structural connections and circuits of the brain and their relationships to behavior by acquiring high quality magnetic resonance images. We used diffusion MRI data from the Human Connectome Project Development (HCP-D) study, the Human Connectome Project Young Adult (HCP-YA) study, and the Human Connectome Project Aging (HCP-A) study. HCP-D was composed of 636 subjects with diffusion data, with age ranges between 5-21. HCP-YA was composed of 1065 subjects with diffusion data, with age ranges between 21-35. HCP-A was composed of 720 subjects with diffusion data, with age ranges between 35-100. Thus, the pooled dataset comprised 2421 participants and spanned the ages of 5 and 100.

The diffusion MRI acquisitions were slightly different for each dataset and tailored towards the populations under investigation. For HCP-D and HCP-A, a multi-shell diffusion scheme was used, with b-values of 1500 and 3000 s/mm^2^, sampled with 93 and 92 directions, respectively. The in-plane resolution was 1.5 mm, with a slice thickness of 1.5 mm. Susceptibility, motion, and eddy current corrections were performed using TOPUP from the Tiny FSL package (http://github.com/frankyeh/TinyFSL), a re-compiled version of FSL TOPUP (FMRIB, Oxford) with multi-thread support. The corrections were conducted through the integrated interface in DSI Studio’s (“Chen” release). For HCP-YA, the minimally-preprocessed data (Glasser et al., 2013) from Human Connectome Projects (Q1-Q4 release, 2015) were acquired at Washington University in Saint Louis and the University of Minnesota (Van Essen et al., 2012) using a multi-shell diffusion scheme, with b-values of 1000, 2000, and 3000 s/mm^2^, sampled with 90 directions each. The in-plane resolution was 1.25mm, with a slice thickness of 1.25mm. Susceptibility, motion, and eddy current corrections were performed as part of the HCP minimal preprocessing pipeline, which also used FSL software TOPUP and EDDY algorithms.

### Superficial White Matter Tractography

For every subject, superficial white matter tractography was performed using methodology similar to (Shastin et al., 2021) using MRtrix software (Tournier et al., 2019). **Figure 1** shows the reconstruction, tractography, and segmentation pipeline. Briefly, all data were up-sampled to 1mm isotropic resolution (using *mrgrid* command), and multi-shell multi-tissue constrained spherical deconvolution (*dwi2fod*) (Jeurissen et al., 2014) was performed to derive estimates of the orientation distributions of white matter fibers (**Figure 1**, left). Tractography was performed using the second-order integration probabilistic algorithm (*tckgen*) (Tournier et al., 2010) to generate 2 million streamlines seeded from the white/gray matter boundary, with a maximum length of 50mm, using anatomical constraints to ensure gray matter to gray matter connections (Smith et al., 2012), in combination with a number of deep gray matter and deep white matter exclusion zones (Shastin et al., 2021). This pipeline has been shown to yield dense systems of fibers directly below the cortical sheet (Shastin et al., 2021) (**Figure 1**, middle). Finally, a tract density map was created (*tckmap*) (Calamante et al., 2011) at 250um isotropic resolution and thresholded at an empirically derived value of 5 streamlines per voxel (see Discussion on Limitations), resulting in the high-resolution map of the locations of SWM systems (**Figure 1**, right).

**Figure 1.**
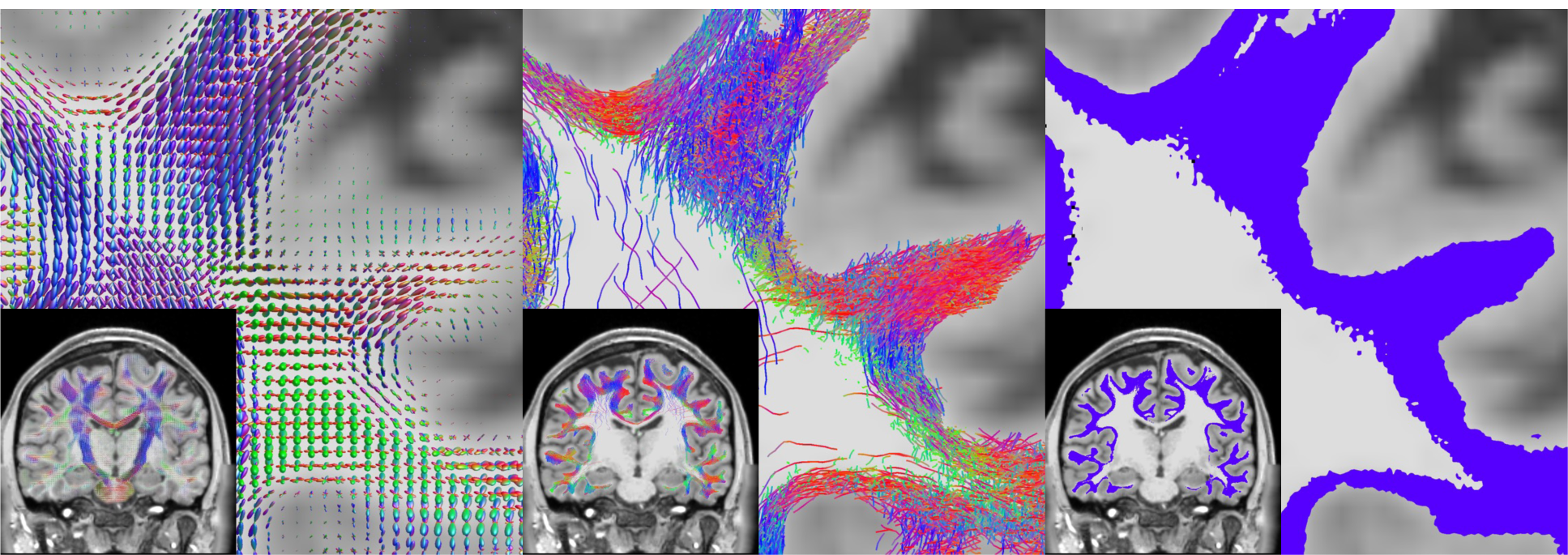
Superficial white matter tractography was performed using methodology described in (Shastin et al., 2021) in order to derive SWM segmentations. This included multi-shell multi-tissue spherical deconvolution (Jeurissen et al., 2014) to derive fiber orientation distributions (left), anatomically informed probabilistic tractography (Tournier et al., 2010) and filtering (middle), and tract segmentation (right).

We performed analyses both using an ROI-based approach and a vertex-based approach with vertices spanning the entire white/gray matter boundary. To this end, the T1-weighted images were analyzed with Freesurfer (Fischl, 2012), and results were transformed to diffusion MRI space with ANTs (Avants et al., 2011). From this, we used both the surface file representing the white/gray matter boundary and the Destrieux atlas (Destrieux et al., 2010) parcellation, resulting in 164 neocortical labels for ROI-based analysis.

### Features

Two features of the SWM were extracted for this study. First, total SWM volume was calculated based on the SWM segmentation by multiplying the number of voxels within the segmentation by the total voxel volume. SWM volume is a single scalar measure per subject. Second, SWM thickness was calculated for every vertex of the white/gray matter boundary. This was done by querying from the vertex *into the white matter* (and orthogonal to the white/gray matter boundary) until U-fibers were no longer detected. This was done in increments of 250*μ* m until the SWM segmentation was no longer encountered. Once the end of the SWM was reached, two options were possible. If the query point was now within the deep white matter, the distance was entered directly as SWM thickness. If the query point reached gray matter, we divided the traveled distance by two as the measure of SWM thickness. This situation arose primarily along sulcal walls, where SWM of neighboring sulci is not distinguishable within the gyral blade. At the end of this procedure, each vertex of the white/gray matter boundary mesh had an associated SWM thickness, and the average value of each cortical region of interest could be quantified for analysis.

Four cortical features were additionally extracted directly from Freesurfer results: (1) cortical thickness, (2) curvature (sulci have positive curvature, gyri negative, with sharper curvature indicated by higher absolute value), (3) the Jacobian of white matter (which computes how much the white matter surface needs to be distorted to register to the spherical atlas), and (4) sulcal depth (which describes distance from the mid-surface between gyri/sulci; sulci have positive values and gyri have negative values). Note that these features follow FreeSurfer conventions and definitions.

### Analysis

Volumetry across the lifespan was analyzed using covariate-adjusted restricted cubic spline regression (C-RCS) (Huo et al., 2016), a flexible approach to model nonlinear relationships between variables. Here, we used knots at 12, 19, 30, 75, and 90 years of age, based on previously identified developmental shifts in volumetry (Hedman et al., 2012), with 5 knots being common as a compromise between flexibility and overfitting (Stone, 1986). From this, the percent change per year was calculated as a measure of change divided by original value times 100. Results were derived binned for age groups pre-determined based on those in the HCP datasets, development (D) (age 5-21), young adult (YA) (21-35), and aging (A) (35-100). To test for a relationship between U-fiber thickness and features of the cortex across a population for each vertex, using the cortical features of all participants as the independent variable and U-fiber thickness as the dependent variable, we fit a robust Pearson’s correlation coefficient using skipped-correlations (Wilcox, 2004) and depicted this cross-sectional correlation coefficient across the brain. We note that similar results were obtained using traditional Pearson’s correlation coefficient (data not shown).

## Results

### What does the SWM thickness look like?

We begin by examining U-fiber thickness in individual subjects. **Figure 2** shows the U-fiber segmentation (top), the ROI-based SWM thickness (middle), and the vertex-based SWM thickness (bottom) for 3 randomly selected subjects in the HCP-D, HCP-YA, and HCP-A cohorts. Qualitatively, SWM runs immediately below and adjacent to the cortex in the expected geometries and locations described in the literature (Guevara et al., 2020; Oishi et al., 2008; Phillips et al., 2013; Zhang et al., 2018). Averaged across regions, SWM thickness largely falls between 2-4.5mm throughout the brain, whereas vertex-based measures show a much wider range, with some vertices indicating SWM thickness >10mm. Both vertex-based and ROI-based visualizations indicate moderate hemispheric symmetry on an individual basis, along with general patterns of high SWM thickness in the frontal lobe, in particular the precentral gyrus and superior frontal cortex, with lower thickness in the temporal and parietal lobes.

**Figure 2.**
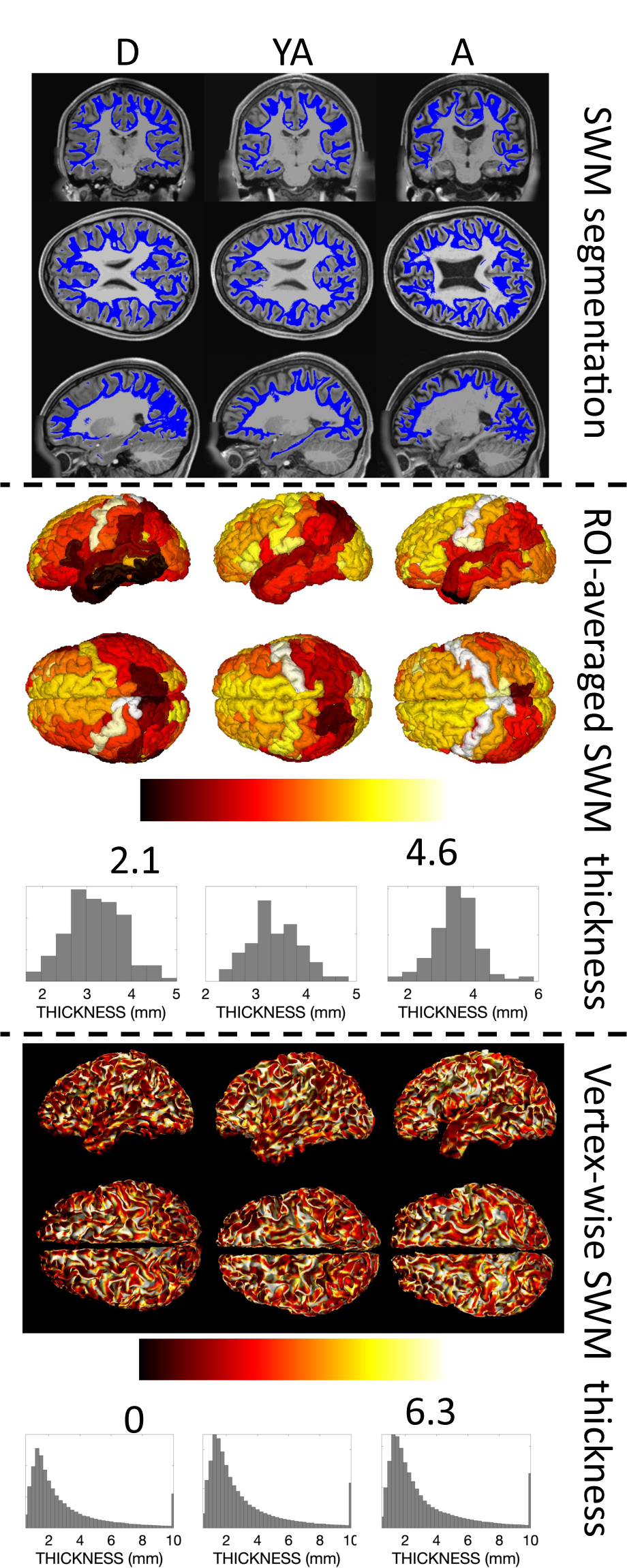
Qualitative differences in SWM thickness are visible across the brain and across the lifespan in individual subjects. U-fiber segmentation (top), ROI-average SWM thickness (middle), and vertex-wise SWM thickness (bottom) are shown for three randomly selected subjects from the development (D), young adult (YA), and aging (A) cohorts.

We next ask what the U-fiber thickness looks like averaged across the population, and visualize this for the D, YA, and A cohorts separately in **Figure 3**. Here, hemispheric symmetry is much more apparent. Average values again fall largely between 2-4mm with a wider range in the finely grained vertex-averaged values. In the ROI-averaged image, similar spatial patterns of thickness are observed across lobes and gyri, and temporal patterns qualitatively suggest an increase between children and young adults, with small changes into aging that vary slightly by location. In the vertex analysis, the gyral crowns result in a greater thickness measurement than walls or fundi, which is likely a result of the chosen quantification procedure. Notably, throughout both gyri and sulci, higher SWM thickness values are observed again in the frontal lobes and throughout the occipital lobes.

**Figure 3.**
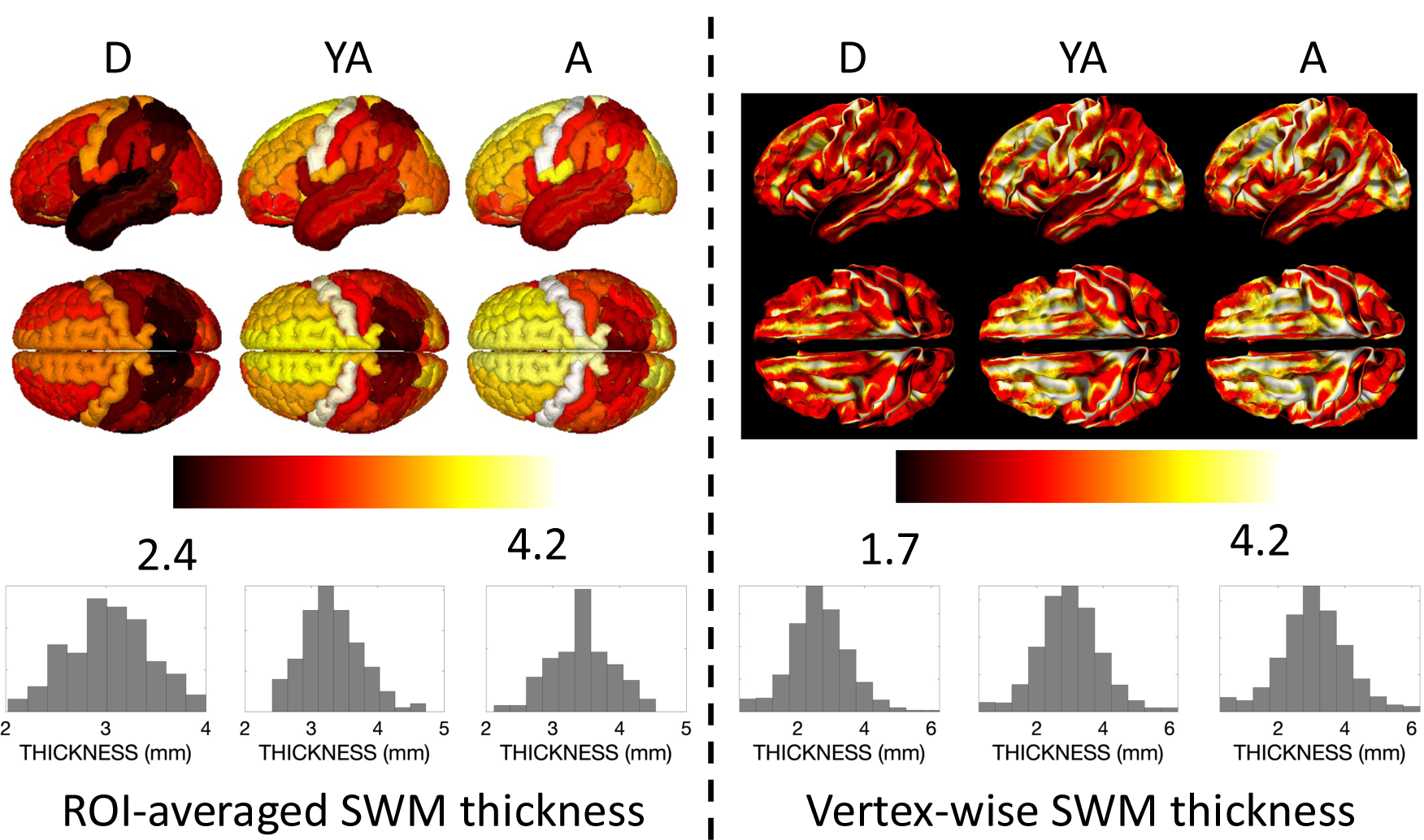
Qualitative differences in SWM thickness are visible across the brain and across the lifespan for the population-averaged subjects for each cohort. The average ROI-based SWM thickness (left) and vertex-based SWM thickness (right) is shown for each of the development (D), young adult (YA), and aging (A) cohorts.

### How does SWM volume change as a function of age?

Changes in volume for the whole brain, gray matter, white matter, and SWM are shown in **Figure 4** (top). Total brain size decreases with age beginning in childhood, (note our cohort starts at 5 years of age) (Bethlehem et al., 2022; Hedman et al., 2012; Scahill et al., 2003; Takao et al., 2012; Taki et al., 2011). Similarly, gray matter volume also decreases with age, with gray matter decreasing faster throughout development and early young adulthood. Also as previously described in the literature, white matter volume increases throughout adolescence, reaching peak volume between 20-35 years of age (Bethlehem et al., 2022; Lebel et al., 2012). Here, for the first time, we find that U-fiber volume shows a similar trend as overall white matter volume, peaking at a similar time in adolescence, leveling off and eventually decreasing throughout adulthood.

**Figure 4.**
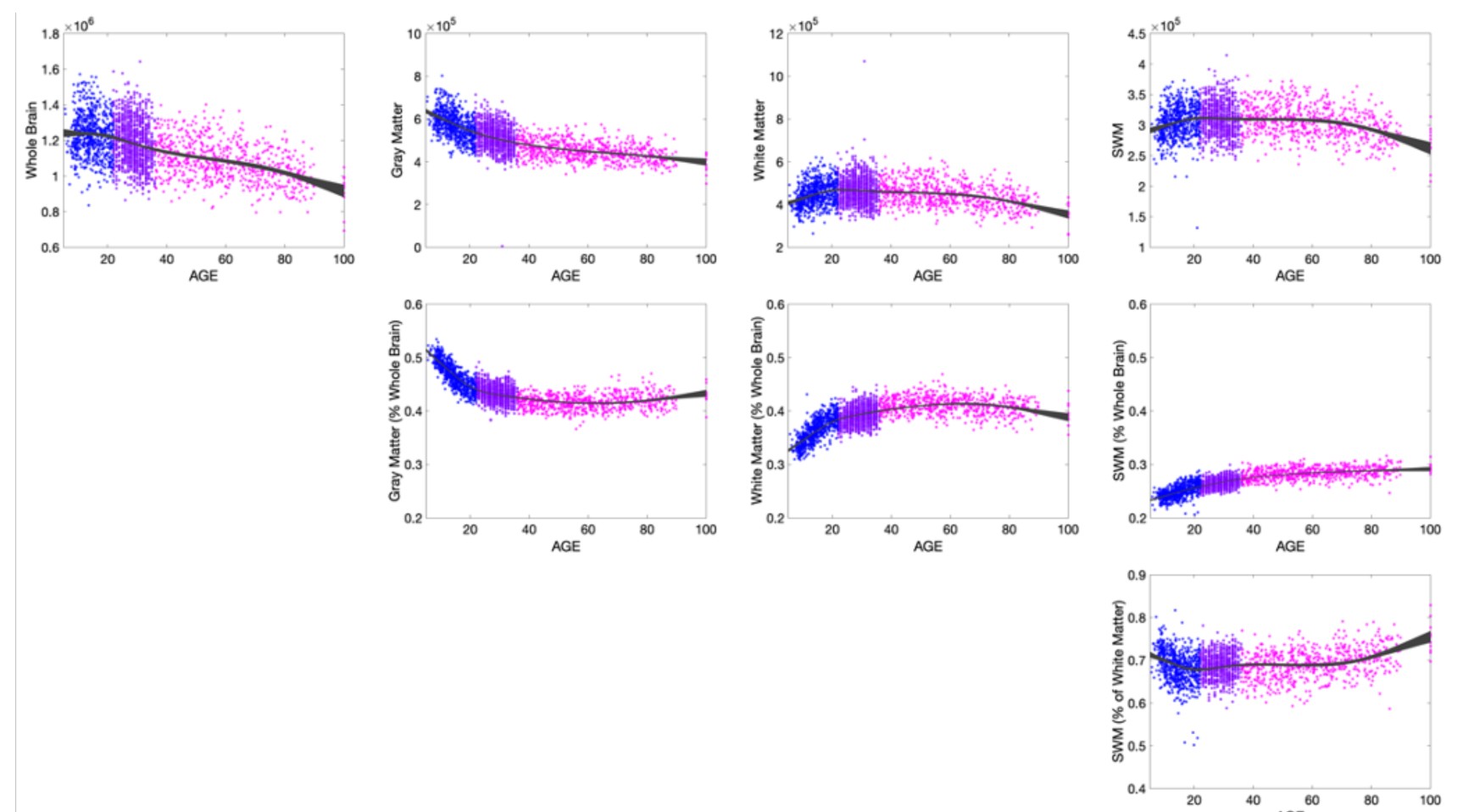
SWM shows volumetric trajectory with age that is unique from total white matter and total gray matter volumes. (Row 1) Volumetric changes in total brain volume, gray matter, white matter, and SWM volumes. (Row 2) Changes relative to whole brain volume. (Row 3) Changes in SWM relative to total white matter volume. Each dataset is shown with a different marker color.

The tissue changes relative to whole brain volume show intuitive changes (**Figure 4**, middle) (Hedman et al., 2012; Takao et al., 2012; Taki et al., 2011; Terribilli et al., 2011). The % gray matter decreases initially until young adulthood then remains relatively constant throughout life, while white matter again increases, reaching a peak value relative to the whole brain throughout mid-life, then decreases with age. However, the % SWM indicates continual increases with age.

Finally, the percentage of white matter occupied by SWM is plotted (**Figure 4**; bottom). U-fibers occupy a majority of the white matter volume, with the % occupied showing a slight decrease during development, and a small but gradual increase with age throughout the lifespan.

### How does SWM volume thickness change as a function of age?

**Figure 5** visualizes the percent change per year in SWM thickness, averaged across the cohorts for D (5-21 years old), YA (21-35), and A (35-100) shown for both ROI-based and vertex-based thickness measures. The most pronounced increase in SWM thickness is observed from age 5-21, with prominent changes seen in the frontal, temporoparietal, and superior temporal gyri. This change is observed throughout the gyral crowns, sulcal walls, and fundi, but is most conspicuous at the gyral crowns. SWM thickness shows similar patterns of increases—albeit of a smaller magnitude—from 21-35, and levels off or decreases thereafter.

**Figure 5.**
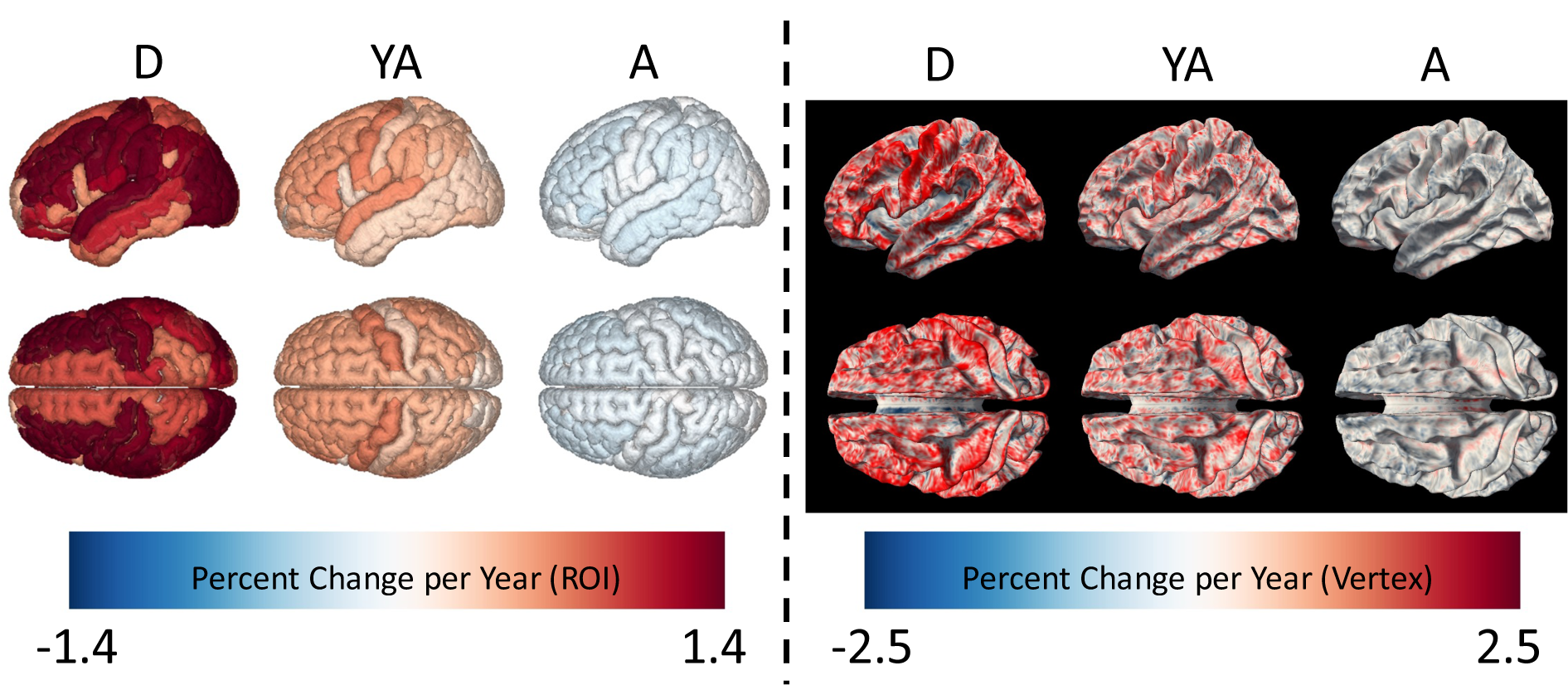
SWM thickness changes throughout the lifespan, with change dependent on location. Left shows ROI-based analysis colored by percent change per year for D, YA, and A cohorts. Right shows the vertex-based analysis colored by percent change per year for the same cohorts.

### Is there a relationship between SWM thickness and key cortical features?

To investigate the relationships between U-fiber thickness and important cortical features (or Freesurfer metrics computed on the cortical surface) – thickness, curvature, Jacobian of the white matter, and sulcal depth – we assessed relationships between features across individuals for every vertex of the surface mesh and visualized the results as cross-sectional correlation coefficients. There was a moderate correlation between cortical thickness and U-fiber thickness throughout the brain, in both gyri and sulci (**Figure 6**), with a correlation coefficient typically greater than 0.2. Thus, as cortical thickness increases, so does U-fiber thickness. Results for curvatures show unique patterns, with positive values in the prefrontal, temporal, and occipital lobes, and negative correlations in the pre and post central gyri and superior temporal gyri. The Jacobian of the white matter shows weak, but positive, correlations with U-fiber thickness, with correlations localized to the gyral crowns, indicating that areas which required larger deformations during transformation to a standard space also have larger U-fiber thicknesses. Finally, sulcal depth shows both positive and negative correlations, with patterns similar to that of curvature.

**Figure 6.**
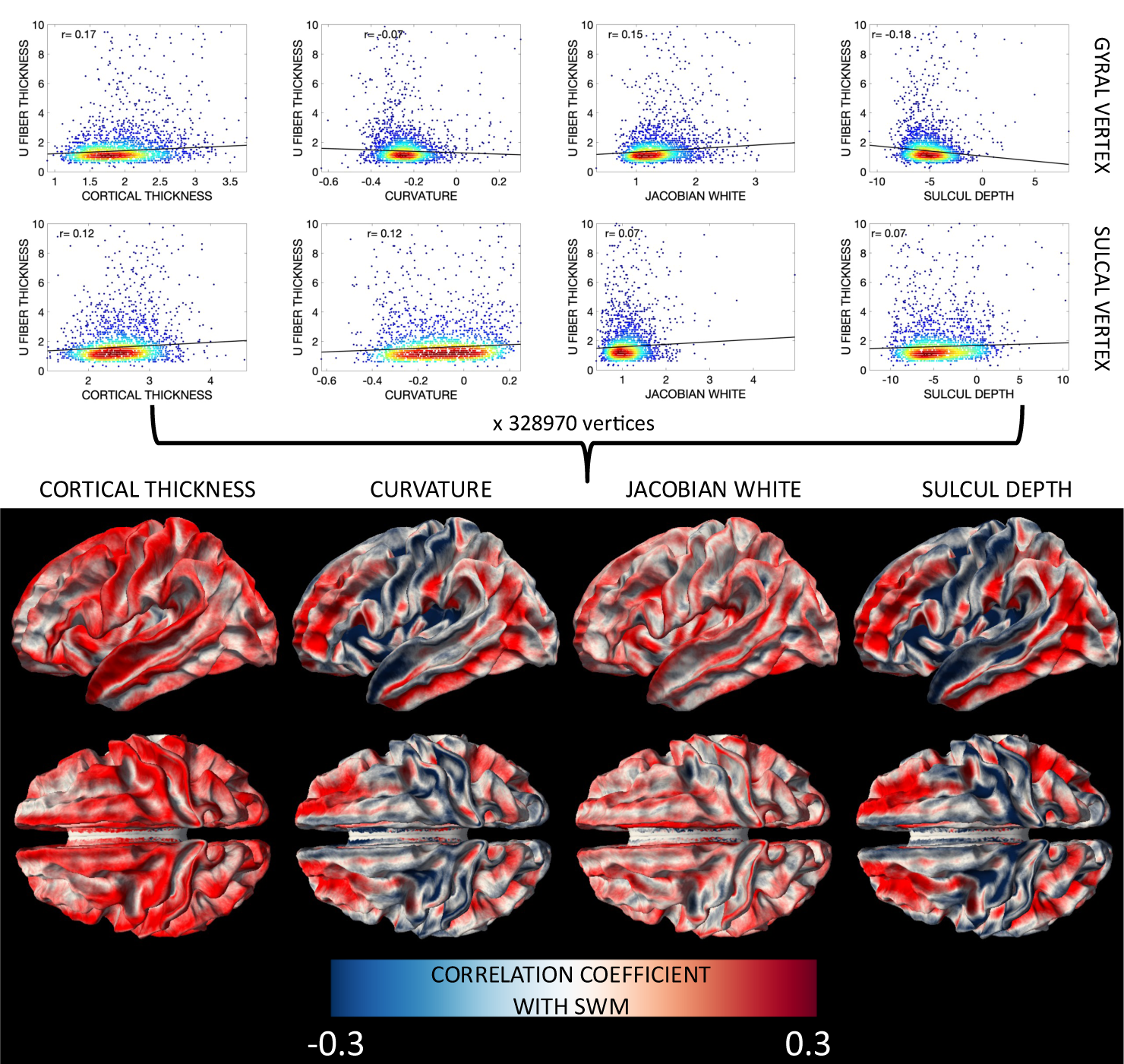
U-fiber thickness shows cross-sectional relationships with FreeSurfer-defined features of the cortex (cortical thickness, curvature, Jacobian of the white matter, and sulcal depth) that vary based on brain location. Here, for every vertex, a robust correlation coefficient between U-fiber thickness and cortical feature across the population was calculated (example plots for one gyral vertex and one sulcal vertex shown in top row) and displayed on the white/gray matter boundary for all vertices (bottom row).

## Discussion

Using a large, high quality, diffusion MRI dataset, along with innovations in tractography and quantification, we characterized SWM in the human brain across most of the lifespan. Our main findings are that (1) SWM volumetric trajectories with age are distinct from gray matter and total white matter trajectories (2) SWM thickness shows patterns robust across individuals and across the population at the region-level and vertex-level, (3) SWM shows spatially-varying nonlinear changes across the lifespan, and (4) SWM thickness is associated with various features of the corresponding cortex. These findings are discussed in detail below.

### SWM volume and age

The brain is known to undergo significant changes during the lifespan. Studies of brain volume, white and gray matter tissue volume (Fan et al., 2019; Fjell and Walhovd, 2010; Hedman et al., 2012; Lebel et al., 2012; Pfefferbaum et al., 1992; Scahill et al., 2003; Steffener, 2021), cortical thickness (Dominguez et al., 2021; Frangou et al., 2022; Habeck et al., 2020; Hou et al., 2021), microstructure measures (Beck et al., 2021; Fan et al., 2019; Lebel et al., 2012; Storsve et al., 2016), and myelination (Grydeland et al., 2019) have described patterns of neuroanatomical variation that provide insight into the biological sequelae of development and aging. For example, well characterized waves of brain growth occur, with gray matter volume increasing until middle childhood (~ 6 years), followed by volume decreases from young adulthood and into late adulthood, while white matter reaches peak volume in adulthood (20-40 years), again leveling off and decreasing into late adulthood, with both tissues experiencing accelerated atrophy during late adulthood (Bethlehem et al., 2022; Hedman et al., 2012; Lebel et al., 2012; Scahill et al., 2003). Within each tissue type, changes are heterogenous, with different growth trajectories in different cortical regions and white matter pathways (Lebel et al., 2012; Lebel et al., 2008; Schilling et al., 2022a), and different associations with axonal myelination (Grydeland et al., 2019) and densities (Beck et al., 2021; Cox et al., 2016; Giorgio et al., 2010; Schilling et al., 2022a). These studies have driven hypotheses that aim to relate structure and function and determine structural mediators or cognition (Armstrong et al., 2020; Frangou et al., 2022; Winter et al., 2021), or may serve as a benchmark of normative trajectories that might be used to reveal patterns of abnormal variation or vulnerability in disease and disorder (Bethlehem et al., 2022; Fjell and Walhovd, 2010; Shafer et al., 2022).

However, similar studies of SWM have not previously been performed, and measurements of SWM volume and thickness have not been documented. Here, we find that SWM volume trends closely follow those of the total white matter, increasing in volume until young-adulthood during which it levels off and decreases, followed by accelerated decline into late-adulthood. Notably, the relative fraction of total brain volume of these SWM systems increases continually with age, and consequently takes up a larger proportion of total white matter volume, which decreases with respect to total brain volume. While this could be an artifact of processing and tractography (see Limitations, below), it could also indicate relative preservation of these heavily myelinated systems in older age. The fact that our tractography-derived volume fraction estimates (~70%) are well in line with existing literature (>60%) serves as an indirect validation of this quantification. A future study could investigate the sensitivity of these U-fiber metrics to changes in cognition or mobility as a biomarker for abnormal degeneration, and compare the results against traditional cortical or white matter phenotypes.

### U-fiber thickness

We have additionally shown that U-fiber thickness shows unique patterns across the brain, in both individuals and across the population. Thicker SWM values tend to be observed in the frontal and occipital lobes, most prominently in precentral gyri and superior frontal gyri of both hemispheres. This is visible both in ROI-averaged and vertex-based maps, although vertex-based maps suggest large variations within a gyrus. While there is noticeable variation of SWM thickness, most of the brain, for most ages, falls within 2.5-3.5mm (ranging from <1mm to >10mm at some gyral crowns), roughly the same as cortical thickness measures which range between 1-5mm with an average of around 2.5mm (Fischl and Dale, 2000). This estimate is thicker than that provided by susceptibility weighting with estimates of 0.5-2mm thick (Kirilina et al., 2020), which could represent an overestimation due to larger diffusion MRI voxels, or an underestimation from susceptibility, which is based on only the densest band of T2 relaxometry measures. Again, our SWM volumetry measures of ~70% of total white matter volume are in line with existing literature of >60% based on histology (Schüz et al., 2002).

We found that SWM thickness increases throughout the brain across childhood and adolescence, changing at a rate of 1-1.5% per year. The greatest increases occur in the primary motor and sensory areas, with large changes distributed throughout the frontal and occipital lobes, and superior temporal gyri. This perfectly matches the most heavily myelinated areas of the cortex (as derived from T1-weighted over T2-weighted ratios (Glasser and Van Essen, 2011; Grydeland et al., 2019), regions of greatest myelin maturation (Grydeland et al., 2019) (see Figure 1 and Figure 3 from (Grydeland et al., 2019)), and also the areas that show the greatest decline in cortical thickness during the same age period (see Figure 3 from (Frangou et al., 2022)). While all MRI studies may be susceptible to a partial volume effect between cortex and SWM, this suggests a strong temporal relationship between these regions, with increasing myelination, increasing SWM volume, and decreasing cortical thickness. The rates decrease during adulthood; however, the same regions stand out as increasing in thickness with age. These results align with previous studies of the long-range association and projection pathways, where fronto-temporal connections experienced both prolonged development (peaking between 20-40 years of age) and late decline. Thus, there is also a complex interplay between the short- and long-range fibers with similar terminal areas. Finally, the rates decline in late aging, although at a much smaller magnitude. However, it is these declines that should be studied, both in terms of volumetric and microstructure-based features (Phillips et al., 2013; Phillips et al., 2016; Schilling et al., 2022b)), as they relate to normal and abnormal aging.

### Relationship with gray matter

Finally, we show that U-fiber thickness has unique relationships with features of the cortex. For example, thicker U-fibers occur in regions of greater cortical thickness. This phenomenon has not been described previously, but reveals an intuitive relationship whereby subjects with greater cortical thickness in a given region also have a larger number of short fibers in this region. The results of positive relationships between U-fibers thickness and the FreeSurfer derived Jacobian of the white matter serves as a validation that larger SWM requires greater white matter distortion to match the template surface. Finally, curvature and sulcal depth (which are intimately related in that sulci have positive depth and positive curvature, while gyral crowns have negative depth and negative curvature) show unique patterns that vary across the brain. In the precentral and postcentral gyri and superior temporal gyrus, U-fiber thicknesses are negatively correlated with both (i.e., subjects with greater curvature have thinner SWM), whereas the rest of the brain has positive correlations with both (i.e., subjects with less curvature have thicker SWM). Thus, further investigation is warranted to determine whether this mixed pattern of positive and negative correlations changes with age and whether it is related to gyral complexity or spatial constraints during development – or whether it may be a bias in methodology (see below).

### Limitations

Several limitations must be acknowledged. First, tractography faces a number of hurdles, such as partial volume effects with other tissue types (Rheault et al., 2020), limitations in resolving crossing fiber populations (Jeurissen et al., 2013; Schilling et al., 2017), particularly at the cortical interface (Reveley et al., 2015), false positive connections (Maier-Hein et al., 2017) due to bottleneck regions in the brain (Girard et al., 2020; Schilling et al., 2021b), and gyral biases (Schilling et al., 2018; St-Onge et al., 2018). However, care was taken to use anatomical constraints (Girard et al., 2014), in combination with surface-based seeding, exclusion regions, and high angular resolution tractography following validated methodology (Shastin et al., 2021). Second, the relatively coarse resolution of diffusion MRI may result in an overestimation of volume and thickness, which is moderated in part by high resolution tract density imaging and subsequent thresholding which gives sub-voxel resolution of fiber trajectories (Calamante et al., 2011; Calamante et al., 2012), although validation against human histological data is needed. Third, thickness quantification may be overly simplistic. We have chosen an intuitive measure, simply propagating into white matter until U-fibers are no longer detected. While this works well in sulcal fundi, it is not intuitively quantified along walls or gyri (whereas the cortical thickness is well-defined between two tangential surfaces). We have chosen to divide this value by ½ within sulcal walls that project directly to the neighboring sulcal wall (with the assumption that the U-fiber systems associated with each wall take up approximately half the gyral blade). It is unknown what percentage of fibers within a gyral blade connect to each side of the U-shaped gyrus, and our solution is clearly only an approximation. This will also lead to overestimates at the top of gyral crowns, where the thickness may truly be a measure of sulcal length. It remains to be investigated how U-fibers are associated with crowns and whether thickness should be quantified at these locations. Another bias due to methodology is that increased volume fraction of SWM relative to whole brain volume as a function of age could be an artifact of reduced whole brain volume as a function of age and improved ability of tractography to form short connections, both true and false positive connections. Finally, different diffusion acquisition and experimental settings are known to influence tractography results (Schilling et al., 2021a), and makes direct comparisons across sites challenging (Fortin et al., 2018; Ning et al., 2020; Tax et al., 2019). For this reason, while the lifespan trajectories are fit using cubic splines, we have chosen to visualize results within each cohort separately.

## Conclusion

SWM represents a significantly understudied part of the human brain, despite taking up a large portion of its volume and making up a majority of its structural connections. Using multiple, high-quality, datasets with large sample sizes in combination with methodological advances in tractography we quantified changes in SWM across the lifespan. We identified and characterized changes in U-fiber volume and thickness across the lifespan, describing trajectories that are unique from gray matter and other white matter tissue. SWM thickness varies across the brain, is reproducible across the population, and shows heterogenous changes during development, young adulthood, and aging. Finally, we find that SWM thickness is related to other biological and geometrical features of the brain. Thus, SWM is continuously changing during the lifespan, and studies of these systems, together with gray matter and other white matter features, can provide insight into normal and abnormal development and aging.

## Acknowledgements

This work was supported by the National Science Foundation Career Award #1452485, the National Institutes of Health under award numbers R01EB017230, K01EB032898, and in part by ViSE/VICTR VR3029 and the National Center for Research Resources, Grant UL1 RR024975–01.

